# A Sequence determinant in 3’UTR of mRNAs for Nuclear Retention by Paraspeckles

**DOI:** 10.1101/2020.07.19.206417

**Authors:** Audrey Jacq, Denis Becquet, Bénédicte Boyer, Séverine Guillen, Maria-Montserrat Bello-Goutierrez, Marie-Pierre Blanchard, Claude Villard, Maya Belghazi, Manon Torres, Jean-Louis Franc, Anne-Marie François-Bellan

**Author notes:** Marie-Pierre Blanchard, Montpellier Ressources Imagerie, BioCampus, University of Montpellier, CNRS, INSERM, Montpellier, France. Manon Torres, Charité–Universitätsmedizin Berlin Campus Charité Mitte Chronobiology in the Charité CrossOver (CCO) Virchowweg 6 Charitéplatz 1 10117 Berlin Germany.

## Abstract

Paraspeckles are nuclear membraneless structures composed of a long non-coding RNA, Nuclear-Enriched-Abundant-Transcript-1 and RNA binding proteins, which associate numerous mRNAs. It is therefore believed that their cellular function is to sequester in the nucleus their associated proteins and/or target mRNAs. However, little is known about the molecular determinant in mRNA targets that allow their association to paraspeckles except that inverted repeats of Alu sequences (IRAlu) present in 3’UTR of mRNAs may allow this association. While in a previous study we established the list of paraspeckle target RNAs in a rat pituitary cell line, we didn’t find however, inverted repeated SINEs, the rat equivalent of primate IRAlus in 3’UTR of these RNAs. By developing a candidate gene strategy, we selected a paraspeckle target gene, namely calreticulin mRNA, and we searched for other potential RNA recruitment elements in its 3’UTR, since 3’UTRs usually contain the sequence recognition for nuclear localization. We found a 15-nucleotide sequence, present as a tandem repeat in 3’UTR of this mRNA, which is involved in the nuclear retention by paraspeckles. While being not overrepresented in the 3’UTR of the paraspeckle target RNAs, this recruitment element was present in near 30% of all 3’UTR mRNAs. In addition, since an oligonucleotide containing this sequence binds the paraspeckle protein component HNRNPK, it is proposed that HNRNPK may constitute a bridging protein between paraspeckles and target mRNAs.

## INTRODUCTION

The paraspeckles, discovered in 2002 [2] are nuclear bodies found in almost all of the cultured cell lines and primary cultures from tissues [3], except for embryonic stem cells [4]. They are usually detected as a variable number of discrete dots found in close proximity to nuclear speckles [3]. The structural element of paraspeckles is a long non-coding RNA, nuclear-enriched abundant transcript one (Neat1) which exists under a short and a long transcript generated from the same promoter, previously identified as MENε (Neat1–1) and MENβ (Neat1–2), respectively [5][6]. The long isoform is essential for the formation and maintain of paraspeckles. More than 60 proteins have been identified thus far to be associated with paraspeckles [7], among which 4 RNA-binding proteins, including 3 members of the Drosophila Melanogaster Behavior Human Splicing (DBHS) family proteins (NONO, PSPC1 and SFPQ) and RNA-binding motif protein 14 (RBM14) are considered as core paraspeckle proteins [3][8][9]. Another crucial paraspeckle protein, the heterogeneous nuclear ribonucleoprotein K (HNRNPK), appeared to affect the production of the essential NEAT1-2 isoform by negatively regulating the 3’-end polyadenylation of the NEAT1-1 short isoform [10]. Paraspeckles have been also shown to associate numerous mRNA. It is therefore believed that the cellular function of paraspeckles is to sequester in the nucleus their associated proteins and/or their target mRNAs. In a rat pituitary cell line, the GH4C1 cells, we identified using a Neat1 RNA pull-down procedure [11] followed by RNA sequencing, 3928 RNAs that are paraspeckle’s targets [12]. In these cells, we have shown that the expression of all major components of paraspeckles including Neat1 and core paraspeckle proteins followed a circadian pattern that leads to rhythmic variations in paraspeckle number within these cells [12]. Thanks to their circadian expression pattern and given their functions in mRNA nuclear retention, it was shown that paraspeckles rhythmically retain target RNAs in the nucleus of the cells and that this rhythmic nuclear retention leads to the rhythmic expression of the corresponding genes [12][13]. However, how mRNAs are bound to paraspeckles is largely unknown. In particular, the molecular sequence determinants that allowed mRNAs to be recognized and targeted by paraspeckles are poorly understood even if it has been shown that inverted repeats of Alu sequences (IRAlu) present in 3’UTR of mRNAs allow these mRNAs to associated with paraspeckles [4][8][14]. Accordingly we and others have shown that the insertion of IRAlus in the 3’-UTR of an Egfp reporter gene induced both a nuclear retention of Egfp mRNA and a reduction of EGFP protein expression in the cytoplasm [14][12]. Moreover, endogenous mRNAs containing IRAlus in their 3’-UTRs like Nicolin 1 (NICN1) or Lin 28 have been shown to be retained in the nucleus by paraspeckles [14][15] providing conclusive evidence that the presence of IRAlus in 3’-UTRs of genes allows paraspeckles to sequester mature mRNAs within the nucleus.

In human cells, hundreds of genes contain inverted repeated short interspersed nuclear elements (SINEs) (mainly Alu elements) in their 3’-UTRs. Alu elements are unique to primates and account for almost all of the human SINEs and >10% of the genome. Their abundance leads to the frequent occurrence of IRAlus in gene regions. Unexpectedly however, in 3’-UTR of the 3928 RNAs we identified as Neat1 targets in rat GH4C1 cells, we didn’t find inverted repeated SINEs, the rat equivalent of primate IRAlus (unpublished results). The question whether other sequence determinants than IRAlus in 3’UTR of mRNAs allow them to be paraspeckle’s targets remains then unanswered. To identify potential RNA recruitment elements, we chose to adopt a gene candidate strategy based on the analysis of the properties of the candidate gene 3’-UTR to associate to paraspeckles. To this end, we selected a candidate gene, namely calreticulin (Calr) mRNA, that was previously shown to be both a post-transcriptional cycling transcript in the liver [16] and a mRNA paraspeckle target whose circadian nuclear retention was dependent on Neat1 in our cell line [12]. We showed that the association of Calr mRNA with paraspeckles involved its 3’UTR that contained a 15-nucleotide sequence, found as a tandem repeat. This sequence determinant was shown to be involved in the nuclear retention of the 3’UTR of Calr mRNA by paraspeckles and was further shown to be present in near 30% of the 3’UTR of the 3928 paraspeckle target RNAs. Finally, as an oligonucleotide containing this sequence was shown to bind the paraspeckle protein component HNRNPK, it was accordingly proposed that HNRNPK could constitute a bridging protein between paraspeckles and target mRNAs through involvement of this 15-nucleotide sequence.

## MATERIAL AND METHODS

### Cell line culture and preparation of stably transfected cell lines

GH4C1 cells, a rat pituitary somatolactotroph line, were obtained in 2012 from ATCC (CCL-82.2, lot number: 58945448, Molsheim, France) with certificate analysis and were confirmed to be free of mycoplasma (MycoAlert, Lonza, Levallois-Perret, France). They were grown in HamF10 medium supplemented with 15% horse serum and 2% fetal calf serum. GH4C1 cells were synchronized between themselves by a replacement of fresh medium. For the generation of stable GH4C1 cell lines, cells were transfected with the plasmid constructs expressing a neomycin resistance gene by Lipofectamine 3000 (Invitrogen, Cergy Pontoise, F). Cells were selected with 250 mg/ml G418 (Invitrogen) beginning 48 h after transfection.

### Plasmid constructs

3’UTR of Calr (3’UTR-Calr) mRNA (574 bases) was amplified using classic 3’ RACE by dual PCR from total RNA obtained from GH4C1 cells (see Supplemental Table 1 for primers). Sequence was then inserted in the plasmid pEGFP-C1 (Clontech, Mountain View, CA) at Sma1 site. Using this 3’UTR construction, PCR were performed in order to obtain three overlapping subsequences named C1, C2 and C3 (see green segments in Supplemental Fig. 3). These sequences were inserted using SmaI site in pEGFP-C1. Primers (IDT, Coralville, Iowa, USA) used for these constructions were listed in Supplemental Table 1. All plasmid constructs were verified by sequencing (Genewiz, Leipzig, Germany).

**Table 1:** Mass spectrometry identification of 24 proteins specifically bound to an oligonucleotide containing the 15-nucleotide sequence.

| Gene names | Peptides | Sequence coverage [%] | Score |
| --- | --- | --- | --- |
| Hnrnpa2b1 | 8 | 22,9 | 191,04 |
| Elavl1 | 3 | 10,7 | 33,948 |
| Ncl | 9 | 13,7 | 74,054 |
| Ssb | 2 | 5,3 | 24,37 |
| Apex1 | 5 | 19,6 | 38,015 |
| Actb | 4 | 14,4 | 33,149 |
| Hnrnpk | 8 | 21,8 | 163,5 |
| Eef1a1 | 4 | 8 | 25,485 |
| Hspa8 | 11 | 18,9 | 183,02 |
| Ptbp1 | 11 | 25,8 | 323,31 |
| Fubp1 | 7 | 13,5 | 63,543 |
| Caprin1 | 4 | 7,6 | 69,329 |
| Rpp25 | 3 | 19,1 | 22,532 |
| Fen1 | 2 | 7,1 | 12,445 |
| Ybx3 | 14 | 57,3 | 323,31 |
| Ptbp2 | 7 | 16,9 | 86,032 |
| Hnrnph2 | 2 | 5,8 | 51,4 |
| Hnrnpa3 | 2 | 10 | 31,933 |
| Ilf2 | 4 | 10,2 | 28,089 |
| Khsrp | 4 | 7,9 | 38,154 |
| Ilf3 | 6 | 7,2 | 45,276 |
| Puf60 | 9 | 19,9 | 104,4 |
| Ptbp3 | 3 | 8,2 | 17,253 |
| Hnrnpd | 12 | 37,4 | 215,08 |

**Fig. 1:**
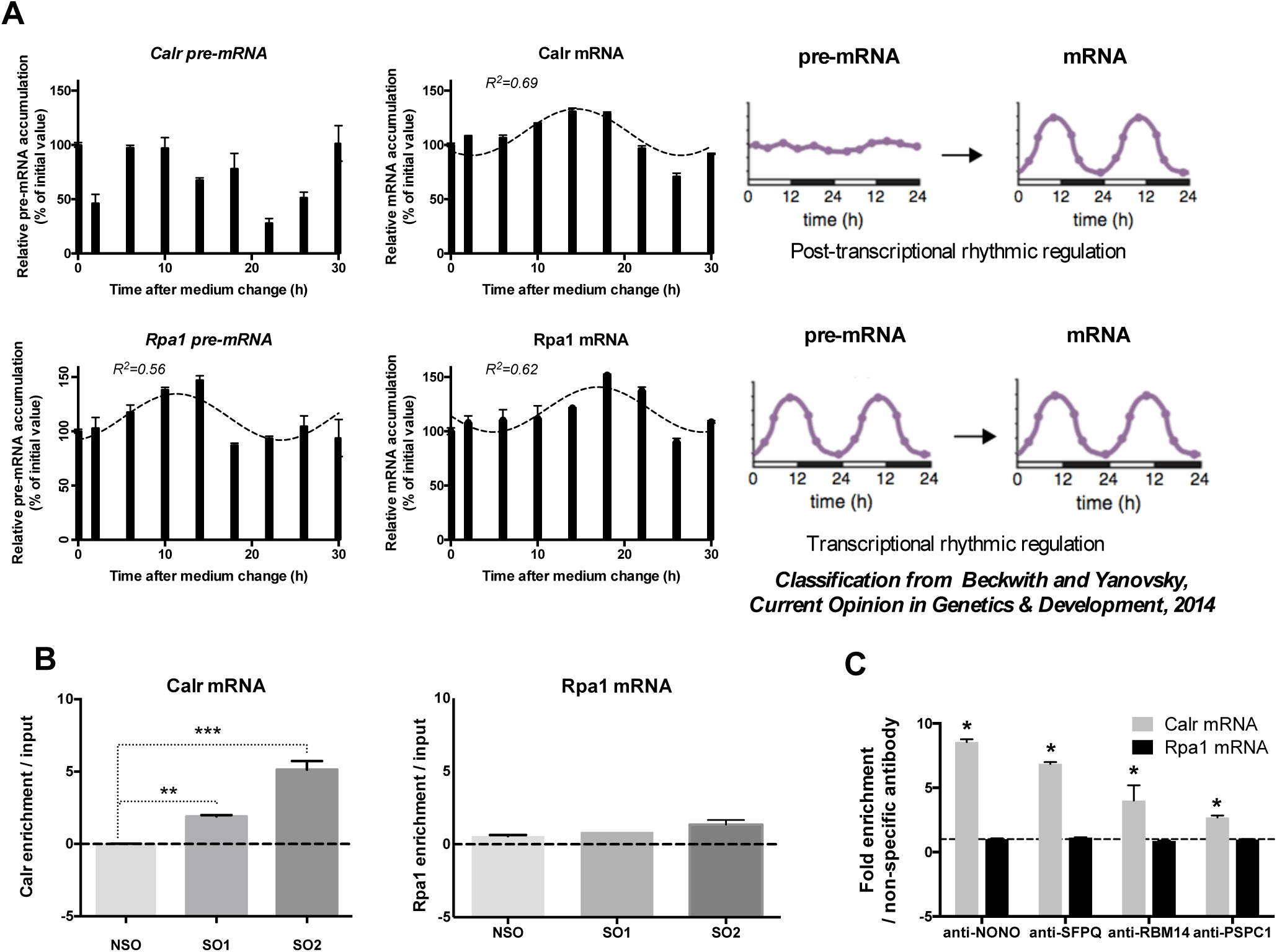
Post-transcriptional control of Calr mRNA circadian expression and association of Calr mRNA with paraspeckle components. **A. Post-transcriptional control of Calr mRNA circadian expression:** The expression of Calr pre-mRNA and Calr mRNA is determined by RT-qPCR over a 30 h time period. Experimental values of Calr pre-mRNA expressed as a percent of the initial value obtained at T=0 (Time of medium change) cannot be adequately fitted (R^2^<0.55) with a non-linear sine wave equation in which the period value is set to 24 h. By contrast, experimental values of Calr mRNA expressed as a percent of the initial value obtained at T=0 can be adequately fitted (R^2^>0.55) with a non-linear sine wave equation in which the period value is set to 24 h. According to the classification from Beckwith and Yanovsky [22] such rhythmic mRNA pattern associated with arrhythmic pre-mRNA pattern, corresponds to post-transcriptional rhythmic regulation (Right Panel). At the opposite the rhythmic expression pattern of Rpa1 pre-mRNA and Rpa1 mRNA give an example of a mRNA submitted to a transcriptional rhythmic regulation according to the classification from Beckwith and Yanovsky [22] (Right Panel). Indeed experimental values of both Rpa1 pre-mRNA and Rpa1 mRNA obtained by RT-qPCR over a 30h time period and expressed as a percent of the initial value obtained at T=0 (Time of medium change) can be adequately fitted (R^2^>0.55) with a non-linear sine wave equation in which the period value is set to 24 h **B. Association of Calr mRNA with Neat1** Enrichment in Calr mRNA relative to input after Neat1 RNA pull-down with two different specific biotinylated oligonucleotides, Specific Oligonucleotide 1 (SO1) and Specific Oligonucleotide 2 (SO2) as compared to a non-specific oligonucleotide (NSO) shows the association of Calr mRNA with Neat1; **p<0.01 ***p<0.001. At the opposite, the lack of enrichment in Rpa1 mRNA relative to input after Neat1 RNA pull-down with SO1 and SO2 compared to NSO attests that Rpa1 is not associated with Neat1 **C. Association of Calr mRNA with four core paraspeckle proteins** Enrichment in Calr mRNA after RNA Immuno-Precipitation (RIP) with antibodies directed against NONO, SFPQ, RBM14 and PSPC1 relative to an irrelevant antibody shows the association of Calr mRNA with the four core paraspeckle proteins; *p<0.05. By contrast, in the same experiments, antibodies directed against NONO, SFPQ, RBM14 and PSPC1 don’t provide an enrichment in Rpa1 mRNA as compared to an irrelevant antibody showing that Rpa1 mRNA is not associated with core paraspeckle proteins.

**Fig. 2:**
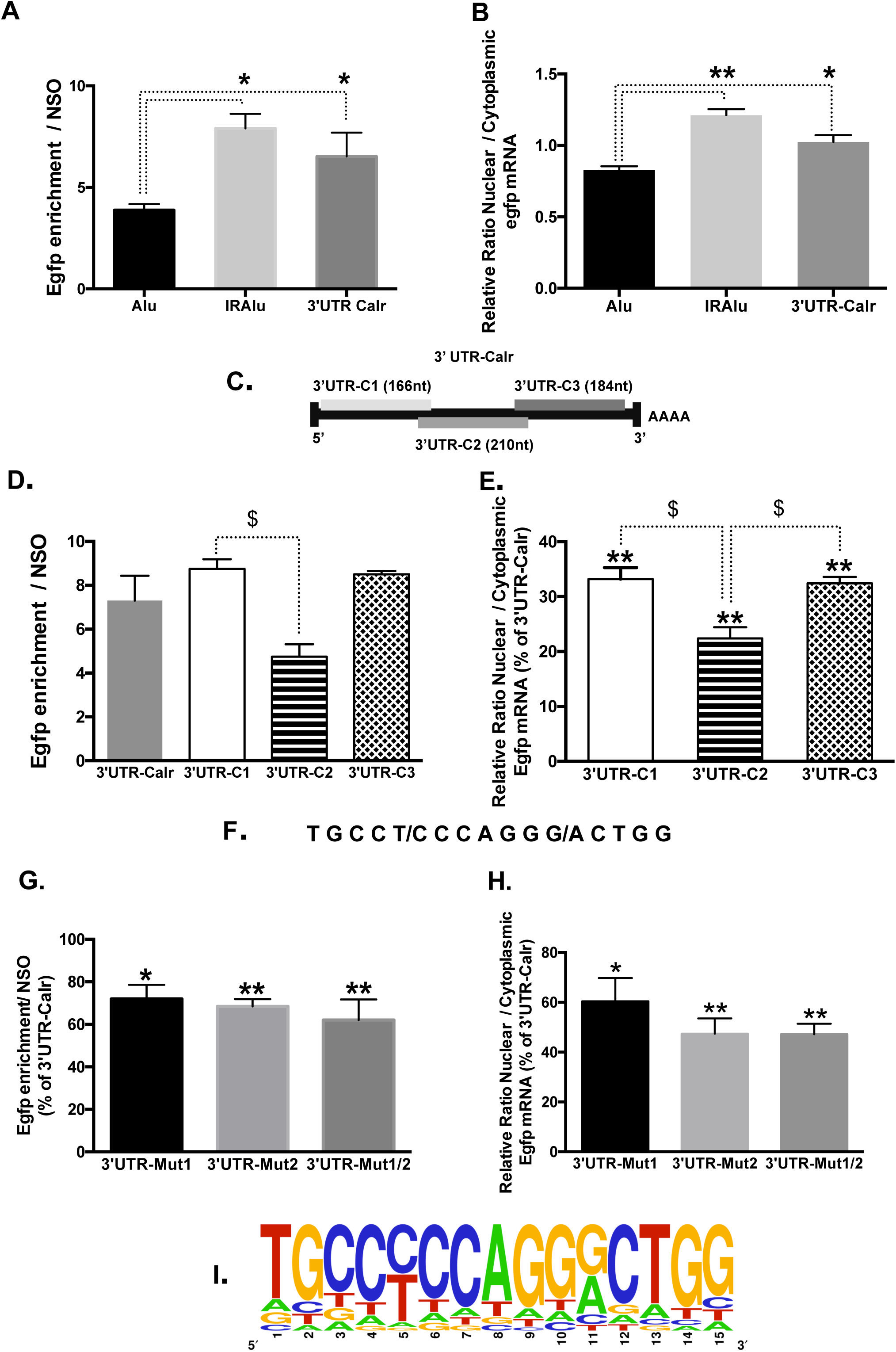
A 15-nucleotide sequence in 3’UTR involved in the nuclear retention of Calr mRNA by paraspeckles: **A-B: Involvement of 3’UTR in Calr mRNA association with paraspeckles** **A**. Enrichment in Egfp mRNA after Neat1 RNA pull-down with Specific biotinylated Oligonucleotide 2 (SO2) relative to Non-Specific Oligonucleotide (NSO). The relative enrichment in Egfp mRNA obtained after RNA pull-down (n=6 for each cell line) is statistically different in IRAlu-egfp and 3’UTR-Calr-egfp versus Alu-egfp cell lines but not different between IRAlu-egfp and 3’UTR-Calr— egfp cell lines * p<0.05 **B**. Nuclear and cytoplasmic Egfp mRNA were quantified by qPCR in Alu-egfp, IRAlu-egfp and 3’UTR-Calr-egfp cell lines and normalized to the relative amount of Gapdh mRNA (n=3 for each cell line). Ratio of nuclear versus cytoplasmic Egfp mRNA levels is statistically higher in IRAlu-egfp and 3’UTR-Calr-egfp cell lines compared to Alu-egfp cell line but not different between IRAlu-egfp and 3’UTR-Calr-egfp cell lines; * p<0.05 **p<0.01. **C-E: Sub-regions of 3’UTR-Calr mRNA associated with paraspeckles** **C**. Schematic representation of the three fragments generated from 3’UTR-Calr mRNA. 3’UTR-Calr mRNA was fragmented in three overlapping parts of 166 (3’UTR-C1), 210 (3’UTR-C2) or 184 (3’UTR-C3) nucleotides, respectively. **D**. Enrichment in Egfp mRNA after Neat1 RNA pull-down with Specific biotinylated Oligonucleotide 2 (SO2) relative to Non-Specific Oligonucleotide (NSO). The relative enrichment in Egfp mRNA obtained after RNA pull-down (n=3 for each cell line) doesn’t differ between the entire 3’UTR-Calr- and 3’UTR-C1- or 3’UTR-C3-cell lines but is statistically different between 3’UTR-C1-versus 3’UTR-C2-cell lines; $ p<0.05. **E**. Ratio of nuclear versus cytoplasmic Egfp mRNA levels obtained in 3’UTR-C1, 3’UTR-C2 and in 3’UTR-C3 cell lines are normalized to the relative amount of Gapdh mRNA (n=3 for each cell line) and expressed as a percent of the ratio in 3’UTR-Calr cell line. The ratio obtained in 3’UTR-C1, 3’UTR-C2 and 3’UTR-C3 cell lines are significantly lower than that obtained in 3’UTR-Calr-egfp cell line; **p<0.01. However, ratio obtained in 3’UTR-C2 is significantly lower that that obtained in 3’UTR-C1 and 3’UTR-C3 cell lines; $ p<0.05. **F-I: Identification of a 15-nucleotide sequence involved in paraspeckle association** **F**. Common sequence of 15 nucleotides found in 3’UTR-C1 and 3’UTR-C3 fragments of the 3’UTR-Calr. **G-H:** The complementary inverse base replaced every odd base in the 15-nucleotide sequence present in position 37-51 (3’UTR-Mut1), in position 466-480 (3’UTR-Mut2) or in both position (3’UTR-Mut1/2). **G**. Enrichment in Egfp mRNA after Neat1 RNA pull-down with Specific biotinylated Oligonucleotide 2 (SO2) relative to Non-Specific Oligonucleotide (NSO) obtained in 3’UTR-Mut1-, 3’UTR-Mut2- and in 3’UTR-Mut1/2-cell lines is significantly reduced compared to that obtained in 3’UTR-Calr cell line; *p<0.05 **p<0.01 **H**. Ratio of nuclear versus cytoplasmic Egfp mRNA levels obtained in 3’UTR-Mut1, 3’UTR-Mut2 and in 3’UTR-Mut1/2 cell lines are normalized to the relative amount of Gapdh mRNA (n=3 for each cell line) and expressed as a percent of the ratio in 3’UTR-Calr cell line. The ratio is significantly reduced in 3’UTR-Mut1, 3’UTR-Mut2 and in 3’UTR-Mut1/2 cell lines compared to the ratio in 3’UTR-Calr cell line. *p<0.05 **p<0.01 **I**. Sequence logo that can be designed from the 1894 sequences found in 3’UTR of mRNA paraspeckle targets.

**Fig. 3:**
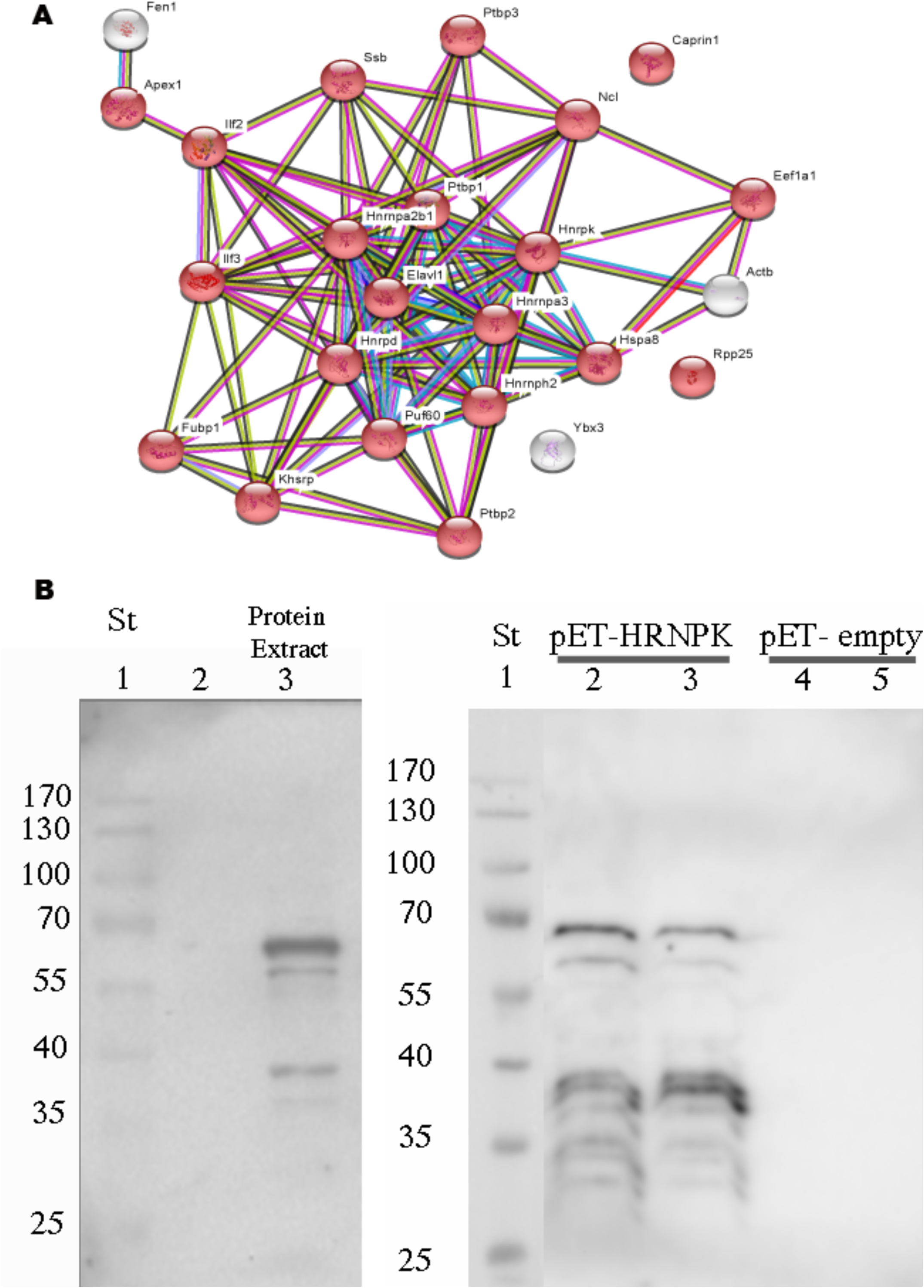
Analysis of proteins associated with the 15-nucleotide sequence identified in 3’UTR-Calr mRNA: Direct binding of HNRNK. **A**. String analysis of the 24 proteins identified by mass spectrometry in eluates from binding to biotinylated RNA containing the 15-nucleotide sequence. Red color: RNA binding protein. **B. Left Panel:** Western blot analysis of HNRNPK in GH4C1 nuclear protein extracts (Lane 3). Lane 1: molecular weight. **Right Panel:** Western blot analysis of the binding of *in vitro* transcribed HNRNPK. Lane 2 and 3: identification of HNRNPK *in vitro* transcribed after RNA-protein pull down using biotinylated RNA specific oligonucleotide containing the 15-nucleotide sequence. Lanes 4 and 5: absence of immunolabeling in the presence of control pET28a lysate. Lane 1: molecular weight

### Mutagenesis

The reporter plasmid pEGFP-C1 containing the entire 3’UTR-Calr mRNA was mutated in the 15-nucleotide sequence present in position 37-51 (Mut1) or in position 466-480 (Mut2) or in both positions (Mut1/2) by inverse PCR using the oligonucleotides listed in Supplemental Table 1. The mutant cDNA sequences were controlled by sequencing (Genewiz).

### RNA expression analysis

From GH4C1 cells, nuclear and cytoplasmic RNA were prepared using the Nucleospin RNA XS kit (Macherey Nagel, Hoerdt, France) and total RNA was prepared using the Nucleospin RNA kit (Macherey Nagel). Nuclear and cytoplasmic RNA isolation was performed using 10 cm cell dishes that were rinsed twice with ice-cold PBS, incubated in 1 ml of ice-cold cell lysis buffer A (10 mmol/L Tris pH 7.4, 3 mmol/L MgCl2, 10 mmol/L NaCl and 0.5% NP-40). Nuclei and cytoplasma were separated by centrifugation (500 g for 5 min). One-sixth of the supernatant was used to prepare cytoplasmic RNA. To obtain pure nuclear RNA, the nuclear pellets were subjected to two additional washes with 1 ml lysis buffer A and were then extracted with Nucleospin RNA XS kit reagent (Macherey Nagel).

The lack of cross contamination between nuclear and cytoplasmic fractions was controlled by western blotting experiments with antibodies directed against a nuclear protein, activating transcription factor 2 (ATF2) (anti-ATF2, sc-187, Santa Cruz Biotechnology, Heidelberg, Germany) on the one hand and a cytoplasmic protein, mitogen-activated protein kinase kinase 1/2, (MEK1/2) (anti-MEK1/2, Cell Signaling, Leiden, The Netherlands) on the other hand.

RNA (500 ng) was then used for cDNA synthesis performed with a High Capacity RNA to cDNA kit (Applied Biosystem, Courtaboeuf, France). Real time PCR was performed on a 7500 fast Real-Time qPCR system (Applied Biosystems) using Fast SYBR Green mix (Applied Biosystems). The sequences of the primers used in qPCR are given in Supplemental Table 1. mRNA accumulation given as the relative ratio of nuclear versus cytoplasmic mRNA was normalized relative to Gapdh mRNA levels.

### RNA immunoprecipitation (RIP) experiments

GH4C1 cells grown in 10 cm dishes were rinsed two times with 5 ml cold phosphate buffer saline (PBS). Cells were then harvested by scraping in ice-cold PBS and transferred to a centrifuge tube. After centrifugation (2500 g for 5 min) cells were pelleted and suspended in 100 μl of Polysome lysis buffer (PLB; 10 mM HEPES, pH 7.0, 0.1M KCl, 5 mM MgCl2, 0.5% NP40, 1 mM DTT, 100 U/ml RNAse OUT and complete protease inhibitor cocktail). After mixing by pipetting up and down, cells were kept on ice for 5 min to allow the hypotonic PLB buffer to swell the cells. The cell lysate was then aliquoted and stored at −80°C.

Cell lysate was centrifuged at 14,000 g for 10 min at 4°C and diluted 1/100 in NET2 buffer (NET2 buffer corresponded to NT2 buffer: 50 mM Tris-HCl, pH 7.4, 150 mM NaCl, 1 mM MgCl2 and 0.05% NP40 added with 1 mM DTT, 20 mM EDTA, 200 U/ml RNAse Out). An aliquot of diluted cell lysate was removed and represented the starting material or ‘input’ which was processed alongside the immunoprecipitation to compare with immunoprecipitated mRNAs at the end. RIP experiments were performed overnight at 4°C on diluted cell lysate with antibodies to NONO (ab45359, Abcam), SFPQ (ab38148, Abcam), PSPC1 (SAB4200068, Sigma-Aldrich, Saint-Quentin Fallavier, France) and RBM14 (ab70636, Abcam) or non-specific rabbit polyclonal antibody (anti-Furin, sc-20801). After incubation was completed, 15 μl of Magna ChIP protein A magnetic beads (16–661, Millipore, Molsheim, France) were added for 1h at 4°C. Beads were washed 6 times with cold NT2 buffer and treated by proteinase K for 30 min at 55°C. RNA eluted from beads was purified using Nucleospin RNA XS (Macherey-Nagel) and processed for cDNA synthesis using a High Capacity RNA to cDNA kit (Applied Biosystems).

### Neat1 RNA pull-down

Neat1 RNA pull-down is a hybridization-based strategy we previously published [11], that uses complementary oligonucleotides to purify Neat1 RNA together with its targets from reversibly cross-linked extracts. Two antisense DNA oligonucleotide probes that target accessible regions of the lncRNA Neat1 were used for Neat1 RNA specific pull-down (SO1 and SO2) and one biotynylated irrelevant probe (NSO) was used for Neat1 RNA non-specific pull-down (Supplemental Table 1). All three probes were biotinylated at the 3’ end.

Briefly, 10 cm cell dishes were incubated in 1 ml of ice-cold cell lysis buffer as described above. Nuclei were scraped and separated by centrifugation (500 g for 5 min). The nuclear pellets were then fixed with 1% paraformaldehyde in PBS for 10 min at room temperature. Crosslinking was then quenched with 1.25 M glycine for 5 min. Cross-linked nuclei were rinsed two times again with PBS and pelleted at 500 g. Nuclear pellets were stored at −80°C. To prepare lysates, nuclear pellets were suspended in lysis buffer (50 mM Tris, pH 7.0, 10 mM EDTA, 1% SDS added with a protease inhibitor cocktail and RNAse-Out), and sonicated using BioruptorPlus (Diagenode, Seraing, Belgium) by 2 pulses of 30 s allowing complete lysate solubilization. RNA was in the size range of 400 to 2000 nucleotides. Nuclear lysates were diluted V/V in hybridization buffer (750 mM NaCl, 1% SDS, 50 mM Tris, pH 7.0, 1 mM EDTA, 15% formamide). The two specific or the non-specific probes (100 pmol) were added to 1.2 ml of diluted lysate, which was mixed by end-to-end rotation at room temperature 4 to 6 h. Streptavidin-magnetic C1 beads (Dynabeads MyOne Streptavidin C1– Invitrogen Life Technologies) were added to hybridization reaction (50 μl per 100 pmol of probes) and the whole reaction was mixed overnight at room temperature. Beads–biotin-probes–RNA adducts were captured by magnets (Invitrogen) and washed five times with a wash buffer (2 SSC, 0.5% SDS). After the last wash, buffer was removed carefully. For RNA elution, beads were suspended in 100μl RNA proteinase K buffer (100 mM NaCl, 10 mM Tris, pH 7.0, 1 mM EDTA, 0.5% SDS) and 100μg proteinase K (Ambion). After incubation at 45°C for 45 min, RNA was isolated using NucleoSpin RNA XS (Macherey-Nagel). Eluted RNA was subject to RT–qPCR for the detection of enriched transcripts.

### *In vitro* transcription/translation

*In vitro* transcription/translation assays were carried out in a S30 T7 HighYield Protein Expression System (Promega). The full lenght Hrnpk cDNA template was cloned from total RNA obtained from GH4C1 cells and transferred to a pET28a *Escherichia coli* T7 expression vector (Novagen, Nottingham, UK). An empty pET28a DNA template was used as control. Reactions were carried out in a total volume of 50μl with 0.5μg plasmid DNA.

### RNA-Protein pull-down

50pmol of a specific 3’ biotin-TEG oligonucleotide RNA probe (Supplemental Table 1) (IDT,) or a non-specific RNA probe (Negative RNA Control [poly(A)_25_ RNA] from the Pierce™ Magnetic RNA-Protein Pull-Down Kit) were incubated for 15-30 min at room temperature under agitation with 50μl of Streptavidin Magnetic Beads. 100 μl of a Master Mix containing 15% glycerol and nuclear GH4C1 protein extracts or proteins translated in vitro (20-200μg) were incubated 30-60 min at 4°C with agitation. After three washes, protein complexes were eluted from RNA by adding 50 μl of Elution Buffer to the beads and incubating 15-30 min at 37°C with agitation. Samples were then loaded onto a NuPAGE 4-12% Bis-Tris gel (Invitrogen) for mass spectrometry analysis or a 10% acrylamide Tris-Glycine gel (Invitrogen) for western blotting.

### MS analysis

Samples were allowed to migrate 5 min at 200 V. The gel was then silver stained and cut into small pieces. Before trypsin digestion, the gel pieces were destained, rinsed and then reduced with dithiothreitol 20mM and alkylated with iodoacetamide 10mM in 100mM NH4HCO3.

Samples were then incubated overnight at 37°C with 12.5 ng/μL trypsin (sequencing grade; Promega) in 25mM NH4HCO3. Gel pieces were then extracted 3 times with 50% acetonitrile 0.1% formic acid and evaporated to dryness using a speed vac.

The dried samples were resuspended in 8μl of 2% acetonitrile and 0.1% formic acid and analyzed by mass spectrometry using a hybrid Q-Orbitrap mass spectrometer (Q-Exactive, Thermo Fisher Scientific, United States) coupled to a nanoliquid chromatography (LC) Dionex RSLC Ultimate 3000 system (Thermo Fisher Scientific, United States). Peptide separation was performed on an Acclaim PepMap RSLC capillary column (75 μm × 15 cm, nanoViper C18, 2 μm, 100 Å) at a flow rate of 300 nl/min. Solvant A was 0.1% formic acid. The analytical gradient was run with various percentages of solvent B (acetonitrile with 0.1% formic acid) in the following manner: (1) 2.5–25% for 57 min, (2) 25–50% for 6 min, (3) 50-90% for 1 min, and (4) 90% for 10 min. MS spectra were acquired at a resolution of 35,000 within a mass range of 400–1,800 m/z. Fragmentation spectra of the 10 most abundant peaks (Top10 method) were acquired with high-energy collision dissociation (HCD).

### Bioinformatics

#### Mass spectrometry data analysis

All mass spectrometry RAW files were uploaded into MaxQuant version 1.5.1.0 (https://maxquant.net/maxquant/, [17]) and searched against a rat SwissProt protein database (December 2018 release). The following parameters were used for the search: trypsin/P enzyme with up to 3 missed cleavages allowed; carbamidomethylation of cysteine was set to fixed modification; with oxidation of methionine, N-terminal protein acetylation set as variable modifications; first search peptide tolerance was set to 20 ppm against a small ‘mouse-first-search’ database for the purpose of mass recalibration and main search was performed at 4.5 ppm; contaminants were included in the search; database was reversed for the purpose of calculating the peptide and protein level false-discover rate (FDR) at 1%; Label Free Quantification (LFQ) minimum-ratio count was set to 1. For statistical comparison a dataset of 3 Specific versus 3 Non-Specific was imported into Perseus software platform version 1.6.2.3 (https://maxquant.net/perseus/, [18]). Contaminants proteins, proteins only identified by site and reverse identifications were filtered out of the dataset. LFQ values were Log2 transformed and samples were filtered with a minimum of 2 for valid values in at least one of the two conditions (Specific and Non-Specific samples).

Results are summarized as a heat Map showing the clustering of the different experiments for the validated proteins.

#### Nucleotide sequence analysis

This analysis was conducted using the bioconductor Biostrings package (v2.52) of R (v 3.6) software.

##### Consensus motif

Local alignment of the 3’UTR-C1 and the 3’UTR-C3 fragments was performed using the pairwiseAlignment function.

##### 3’ UTR sequences

The rat DNA sequences and the coordinates of the 3’UTR were obtained from the Rattus_norvegicus.Rnor_6.0.fa and Rattus_norvegicus.Rnor_6.0.100.gtf in Ensembl, respectively. For each gene, only the sequence of the longest 3’UTR described, when available, was taken into account, leading to a list of 15698 sequences, which was considered as the background file. From this file, a subset of sequences was extracted, those belonging to the 3928 Neat1 RNA targets we previously established (11), leading to a list of 3533 sequences considered as the specific file.

##### Consensus matrix

The specific and the background files were scanned for the TGCCYCCAGGRCTGG motif using the vmatchPattern function, allowing 4 mismatches.

A consensus matrix was generated from the 1894 sequences obtained, using the consensusMatrix function.

A logo was drawn from this matrix, using the seqLogo package (v 1.50) from bioconductor.

### Western-blot analysis

Eluate samples from RNA-protein pull-down experiments were submitted to Western-blot analysis as previously described [19] with polyclonal primary antibodies raised against HNRNPK (1:1000, sc-28380, Santa Cruz Biotechnology).

### RNA-FISH

To detect Neat1 or/and Calr RNA, GH4C1 cells grown on glass coverslips coated with poly-ornithine were fixed in 3.6% formaldehyde. Hybridization was carried out using Custom Stellaris FISH Probes (Biosearch Technologies, Novato, USA). Probes used for dual FISH are Calr probes labeled with Quasar 670 Dye and Neat1 probes labeled with Quasar 570 Dye. Nuclei of the cells were counterstained by Hoechst solution (1μmol/ml).

### Cosinor and statistical analysis

Cosinor and statistical analyses were performed using Prism4 software (GraphPad Software, Inc.).

For cosinor analysis, mean experimental values (± SEM), expressed as a percent of initial value, were fitted using Prism4 software (GraphPad Software, Inc.) by a non-linear sine wave equation: Y = Baseline + Amplitude * sin (Frequency*X + Phase-shift), where Frequency= 2pi/period and period=24h. Goodness-of-fit was quantified using R squared, experimental values being considered well fitted by cosinor regression when the R squared was higher than 0.55. A statistically significant circadian oscillation was considered if the 95% confidence interval for the amplitude did not include the zero value (zero-amplitude test)[20][21].

One-way ANOVA followed by Fisher’s LSD test was used to test for significant differences between groups. Values were considered significantly different for p<0.05 (*), p<0.01 (**) or p<0.001 (***).

## RESULTS

### 1/ Post-transcriptional Calr mRNA circadian expression

According to Menet et al. [16] and Beckwith and Yanovsky [22] the rhythmic pattern of mRNAs is believed to be regulated at the post-transcriptional or transcriptional level depending on whether their corresponding pre-mRNA is arrhythmic or rhythmic (Fig. 1A, Right Panel). While Calr mRNA displayed a circadian expression pattern in GH4C1 cells, Calr pre-mRNA didn’t (Fig. 1A). Experimental Calr mRNA values could indeed be fitted by the following equations: Calr: y= 111.8 – 21.35*sin (0.261x + 0.938) with a R^2^= 0.688. By contrast, Calr pre-mRNA levels could not be significantly fitted by a sine wave equation with a period of 24h (R^2^<0.55) (Fig. 1A). It then appeared that Calr pre-mRNA did not display a rhythmic expression pattern (Fig. 1A), despite high fluctuations of Calr pre-mRNA levels during the time course as previously shown to occur for 80% of pre-mRNA of post-transcriptionally regulated mouse liver cycling transcripts [16]. Accordingly, it appeared that in GH4C1 cells, the rhythmic pattern of Calr mRNA was regulated at a post-transcriptional level. An example of a rhythmic mRNA with circadian transcription was provided with Rpa1 (Replication protein A1), whose values of both mRNA and pre-mRNA could be fitted by non-linear sine wave equations with a period of 24h (Rpa1 pre-mRNA: y= 113.2 + 21.3*sin (0.261x – 1.377) with a R^2^= 0.563 and Rpa1mRNA: y= 120.1 – 20.7*sin (0.261x + 0.287) with a R^2^= 0.617) (Fig. 1A). It then may be concluded that,as expected for a mRNA paraspeckle target, the control of Calr circadian expression pattern took place at a post-transcriptional level, and this was not the case for Rpa1 we didn’t found to be a paraspeckle target in our previous study [12].

### 2/ Association of Calr mRNA with paraspeckles in GH4C1 cells

To provide evidence for the association of Calr mRNA with paraspeckles, we investigated its association with all major components of the nuclear bodies, namely Neat1 and the four major core paraspeckle proteins.

The association of Calr mRNA with Neat1 was investigated using the Neat1 RNA pull down procedure we previously described [12] [11]. In agreement with our previous reports [12] [11], the procedure was shown here to allow a highly significant enrichment in Neat1 RNA by using specific oligonucleotides (SO1 and SO2) compared to non-specific oligonucleotides (NSO) probes (Supplemental Fig. 1A). Furthermore in these experiments, Calr mRNA was shown to be significantly enriched relative to input when SO probes directed against Neat1 were used as compared to a NSO probe, showing clearly that Calr mRNA was associated with Neat1 (Fig. 1B). By contrast, there was no enrichment relative to input in Rpa1 mRNA after use of the two SO probes as compared to NSO (Fig. 1B) showing that Rpa1 was not associate with Neat1 in agreement with our previous results [12]. While Calr mRNA enrichment was statistically significant with the two SO used (SO1 and SO2) compared to NSO, SO2 induced a significantly higher enrichment (Fig 1B); this prompted us to use this oligonucleotide probe in our further experiments.

To investigate the association of Calr mRNA with paraspeckle proteins, we used RNA-immunoprecipitation (RIP) experiments with antibodies directed against the four core paraspeckle proteins, NONO, SFPQ, RBM14 or PSPC1. A 3-to 8.5-fold enrichment in Calr mRNA was obtained with these antibodies as compared to an irrelevant non-specific antibody (Fig. 1C). By contrast, in these experiments, no enrichment in Rpa1 mRNA was obtained with the four specific antibodies showing that Rpa1 mRNA was not associated with core paraspeckle proteins (Fig. 1C). The association of Calr mRNA with the four core protein components of paraspeckles reinforced the view that Calr mRNA bound paraspeckles.

Finally, we performed dual FISH experiments to provide additional morphological indication that Calr mRNA may be associated with paraspeckles visualized by Neat1 RNA staining (Supplemental Fig. 2). As already described, Neat1 RNA staining under confocal laser scanning microscope appeared as regular punctates within the boundaries of the nucleus (Green Supplemental Fig. 2) [12]. By contrast, Calr mRNA staining was mainly localized in the cytoplasm while some punctates also appeared in the nucleus (Red Supplemental Fig. 2). Among these nuclear punctates, some merged with Neat1 RNA staining (Supplemental Fig. 2 arrow) showing that Calr mRNA could be associated with Neat1 RNA. By contrast, some Neat1 RNA stained punctate did not merge with Calr mRNA staining (Supplemental Fig. 2 arrow head).

Taken together, all these results supported the idea that Calr mRNA was closely associated with paraspeckles in GH4C1 cells and validated its choice as a gene candidate to search for molecular mechanism of this association.

### 3/ Involvement of 3’UTR in the binding of Calr mRNA to paraspeckles

Paraspeckles are known to retain in the nucleus RNAs containing duplex structures from inverted repeats of the conserved Alu sequences (IRAlu elements) within their 3’-UTR as shown for Nicolin 1 (NICNI) gene [15]. We used previously generated GH4C1 cell lines stably transfected by constructs in which an IRAlu sequence from the 3’-UTR of NICN1 gene or an Alu element as a control was inserted each between the EGFP cDNA 3’-UTR region and the SV40 polyadenylation signal of the expression vector pEGFP-C1 [23]. To evaluate whether the 3’UTR of Calr mRNA could bind paraspeckle nuclear bodies, the same strategy was used and 3’UTR of Calr mRNA was cloned by PCR, inserted in pEGFP-C1 vector and the construct was stably transfected into GH4C1 cells. Neat1 RNA pull-down experiments were performed in the three cell lines containing constructs with either Alu- or IRAlu element or 3’UTR of Calr mRNA. We verified that Neat1 enrichment obtained with SO2 was significantly higher compared to NSO but not different between the three cell lines (Supplemental Fig. 1B). Consistent with the known ability of IRAlu elements to bind paraspeckles, we found as previously reported [12] that the amounts of Egfp mRNA retrieved after Neat1 RNA pull-down by SO2 compared to NSO were significantly higher in IRAlu compared to Alu-egfp cell line (Fig. 2A). The efficacy of the 3’UTR of Calr mRNA to enrich Egfp mRNA after Neat1 RNA pull-down with SO2 compared to NSO was then compared to that of Alu-or IRAlu-elements. As shown in Fig. 2A, Egfp enrichment after Neat1 RNA pull-down was significantly higher in cell line expressing 3’UTR-Calr as compared to the cell line expressing Alu-containing Egfp mRNA and no statistical difference was obtained between cell lines expressing either 3’UTR-Calr- or IRAlu-containing Egfp mRNA (Fig. 2A). The 3’UTR of Calr mRNA associated as efficiently as IRAlu element with Neat1.

We then performed nuclear and cytoplasmic separation to investigate the influence of 3’UTR Calr mRNA on the nucleo-cytoplasmic distribution of Egfp reporter mRNA. Cross contamination between nuclear and cytoplasmic fractions was ruled out by showing that a nuclear transcription factor, ATF2, and a cytoplasmic protein, MEK1/2, were expressed exclusively in nuclear and cytoplasmic fractions, respectively (data not shown). We showed that both the IRAlu from Nicn1 and the 3’UTR of Calr mRNA caused a significantly greater nuclear retention of the Egfp mRNA when compared with the corresponding Alu element with no significant difference between 3’UTR-Calr and IRAlu element (Fig. 2B). This clearly showed that 3’UTR-Calr sequence retained Egfp mRNA in the nucleus as efficiently as IRAlu sequence did. Taken together, these results were consistent with the involvement of the 3’UTR of Calr mRNA in the binding of Calr mRNA to paraspeckle nuclear bodies.

### 4/ Delineation of sub-regions in 3’UTR-Calr mRNA engaged in paraspeckle binding

To further characterize the sub-regions of the 3’UTR of Calr mRNA that are involved in the binding to paraspeckles, three fragments (3’UTR-C1 (166nt), 3’UTR-C2 (210nt) and 3’UTR-C3 (184nt)) were generated by PCR from the entire 3’UTR of Calr mRNA (Fig. 2C). Each of the three fragments were inserted between the Egfp cDNA 3’-UTR region and the SV40 polyadenylation signal of the expression vector pEGFP-C1 to generate constructs that were then stably transfected into GH4C1 cells. Neat1 RNA pull-down experiments were performed in the cell lines containing constructs with 3’UTR Calr mRNA, 3’UTR-C1, 3’UTR-C2 or 3’UTR-C3. In all these cell lines, Neat1 enrichment obtained with SO2 was significantly higher compared to NSO but there was no statistical difference between the different cell lines (Supplemental Fig. 1C). The efficacy of each fragment was then compared to that of the entire 3’UTR of Calr mRNA to enrich Egfp mRNA after Neat1 RNA pull-down with SO2 versus NSO. Egfp enrichment obtained with 3’UTR-C1 and 3’UTR-C3 fragments did not differ from that obtained with the entire 3’UTR-Calr; by contrast Egfp enrichment obtained with 3’UTR-C2 was reduced and significantly lower than that obtained with 3’UTR-C1 (Fig. 2D). All three fragments were however less able than the entire 3’UTR of Calr mRNA to retain Egfp mRNA in the nucleus; while 3’UTR-C1 and 3’UTR-C3 induced a same retention of about 30% compared to the entire 3’UTR, 3’UTR-C2 was significantly less efficient, inducing a 20% nuclear retention compared to that obtained with the entire 3’UTR (Fig. 2E). It then appeared that while 3’UTR-C1 and 3’UTR-C3 were as efficient as the entire 3’UTR-Calr to induce the binding of Egfp mRNA to paraspeckles, they were not as efficient as the entire 3’UTR-Calr to retain Egfp mRNA in the nucleus. In addition, 3’UTR-C2 was less efficient both to bind the paraspeckles and to retain Egfp mRNA in the nucleus.

### 5/ Involvement of a tandem sequence of 15 nucleotides in 3’UTR-Calr mRNA in the binding to paraspeckles

Through alignment of the oligonucleotide sequences of the fragments C1 and C3 using the Biostring pairwise Alignment function (Smith-Waterman local alignment), a common sequence of 15 bases was identified in these two fragments, with a T or C (Y) in 5^th^ and a G or A (R) in 11^th^ position (Fig. 2F). The 3’UTR of Calr mRNA exhibited then a tandem of this 15-nucleotide sequence (Supplemental Fig 3). To investigate the involvement of these sequences in paraspeckle binding, mutations were introduced in the 15-nucleotide sequence present in position 37-51 (3’UTR-Mut1, Supplemental Fig. 3), or in position 466-480 (3’UTR-Mut2, Supplemental Fig. 3) or in both position (3’UTR-Mut1/2) by replacement of every odd base by its complementary inverse base. 3’UTR of Calr mRNA with Mut1, Mut2 or Mut1/2 were inserted in pEGFP-C1 vector and the different constructs were stably transfected into GH4C1 cells. We verified after Neat1 RNA pull-down experiments in all cell lines that Neat1 enrichment obtained with SO2 was significantly higher compared to NSO but not different between the different cell lines (Supplemental Fig. 1D). Enrichment in Egfp mRNA obtained after these mutations was compared to that obtained with the native entire 3’UTR-Calr mRNA. Mutations in the sequence present in position 37-51 as well as in position 466-480 significantly decreased Egfp mRNA enrichment (Fig. 2G); however the dual mutation of the two sequences didn’t amplify the reduction in Egfp mRNA enrichment (Fig. 2G). While Mut1 and Mut2 significantly reduced the relative ratio of nuclear versus cytoplasmic Egfp mRNA distribution when compared to the native entire 3’UTR-Calr, dual mutation of the two sequences did not display additive or synergic effect (Fig. 2H).

### 6/ Number of occurrences of the 15-nucleotide sequence in the 3’UTR of the Neat1 mRNA targets

The prevalence of the 15-nucleotide sequence identified in the 3’UTR of the Calr mRNA was determined in the 3’UTR of the 3928 Neat1 RNA targets we previously established [12]. Since a same mRNA can display different length of 3’UTR, the list of the Neat1 RNA targets was filtered to retain for each mRNA only the longest 3’UTR referenced. This led to a list of 3533 3’UTR. For this analysis four mismatches were allowed in the 15-nucleotide sequence. In the list of 3533 3’UTR, we found 1894 times this specific sequence included in 1159 mRNA (Table 2). It then appeared that near 30% of the Neat1 RNA targets had at least one identified 15-nucleotide sequence in their 3’UTR, with a mean of 1.6 sequences per gene. However when considering the background defined as total rat RNAs, the 15698 3’UTR retained for the analysis (after filtering for the longest 3’UTR of a single mRNA), 7937 sequences were identified and found to correspond to 4890 mRNA; near 30% of total RNAs displayed then at least one identified 15-nucleotide sequence in their 3’UTR, as Neat1 RNA targets did (Table 2). Accordingly, while the 15-nucleotide sequence we described in the present paper could be involved in the binding of a mRNA to paraspeckles, this sequence was not enriched in the 3’UTR of mRNA that are paraspeckle’s target. Interestingly, it should be mentioned that the same hold also true for IRAlu sequences. It could then be proposed that in the same way than previously reported with IRAlu element, the 15-nucleotide sequence described here contributed to paraspeckle binding but was not sufficient by itself to induce binding to paraspeckle.

**Table 2:** Prevalence of the 15-nucleotide sequence in the 3’UTR of Neat1 RNA targets and in the 3’UTR of all rat RNAs.

|  | Number of<br>3'UTR | Number of<br>sequences | Number of<br>genes | Number of<br>sequences/genes |
| --- | --- | --- | --- | --- |
| **Neat1 RNA**<br>**targets** | 3533 | 1894 | 1159 | 1.6 |
| **Background**<br>**(total RNAs)** | 15698 | 7937 | 4890 | 1.6 |

The important prevalence of this 15-nucleotide sequence in total RNAs prompted us to investigate further its relevance. Since in the 15-nucleotide sequence the 5^th^ and the 11^th^ positions were Y and R, respectively, there were 4 variants of this sequence with each its own number of occurrences. The mean value of those occurrences for the 4 variants was 763 times in the list of the Neat1 RNA targets and 3209 times in total rat RNAs (Supplemental Fig. 4). To evaluate the relevance of these values, we compared them to the number of occurrences of 30 randomly generated sequences of 15 nucleotides with Y or R in the in the 5^th^ and 11^th^ positions, respectively (30 was the minimum size for a statistical relevant sampling for the global population). The mean value of the number of occurrences for the 4 variants of the 30 random sequences was shown to be significantly lower, 363 times and 1608 times in the list of the Neat1 RNA targets and in the total rat RNAs, respectively (Supplemental Fig. 4). It then appeared that the 15-nucleotide sequence we have identified was about twice more frequent than a random sequence of same length in 3’ UTR of both Neat1 RNA targets and total RNAs. Since the prevalence of the 15-nucleotide sequence was statistically relevant, a table of consensus matrix probabilities was therefore built.

The table of consensus matrix probabilities built from the 1894 sequences found in 3’UTR of Neat1 mRNA targets (Supplemental Table 2), allowed the design of a sequence logo (https://rdrr.io/bioc/seqLogo/) (Fig. 2I). The frequency matrix and its graphic representation clearly show that there was a core motif of YCCAGGR (Fig. 2I, Supplemental Table 2).

### 7/ Analysis by mass spectrometry of proteins bound to the 15-nucleotide sequence

To identify proteins able to bind the 15-nucleotide sequence, RNA-protein pull-down was performed using nuclear protein extracts from GH4C1 cells and a biotinylated 30 nucleotides RNA probe corresponding to the 15-nucleotide sequence present in C3 in the 3’UTR-Calr mRNA surrounded in 5’ by 7 nucleotides and in 3’ by 8 nucleotides (Supplemental Table 1). Three replicates using the specific probe and three replicates using a non-specific probe (Negative RNA Control [poly(A)25 RNA] from the PierceTM Magnetic RNA-Protein Pull-Down Kit) were performed. Label free mass spectrometry analysis gave a list of 45 proteins that were identified at least in two biological replicates using the same probe (Specific or Non-Specific) (Supplemental Fig. 5). Finally, 24 from this pool of 45 proteins displayed higher expression in Specific as compared to Non-Specific samples. These 24 proteins were thus considered as specifically bound to the RNA probe containing the 15-nucleotide sequence (Table 1) and were submitted to a String analysis [24]. This analysis showed that 21 of these 24 proteins were RNA-binding proteins (Fig. 3A, red color) and that a dense network connected 21 of each to each other (Fig. 3A). HNRNPK was the only protein in our list that was previously described as a paraspeckle protein component [10].

### 8/ Evidence for the direct binding of the protein HRNPK to the RNA oligonucleotide containing the 15-nucleotide sequence

We tested whether HRNPK could directly bind the RNA oligonucleotide containing the 15-nucleotide sequence identified here. Rattus HNRNPK was then transcribed *in vitro* and incubated with the specific 30 nucleotides RNA probe containing the 15-nucleotide sequence identified in 3’UTR-Calr mRNA. Used as a control, an empty pET28a DNA lysate was also incubated with the specific RNA probe. After elution, samples were processed for western blot with an antibody directed against HRNPK. Western blot analysis of nuclear protein lysates from GH4C1 cells showed that HNRNPK corresponded to a prime band of 65 kDa (Fig. 3B Left panel). As shown in Fig. 3B right panel, a prime band of 65 kDa was also visible in elution fractions from samples where HNRNPK had been transcribed from Hnrnpk-pET28a (lanes 2 and 3) in contrast to samples transcribed with an empty pET28a (lanes 4 and 5). It then appeared that HNRNPK could directly bind the RNA oligonucleotide containing the 15-nucleotide sequence.

## DISCUSSION

Although the physiological functions of paraspeckles remain only partly understood, two known functions of paraspeckles are the sequestration of specific transcription factors and/or RNA-binding proteins and the regulation of the expression of specific transcripts via their retention in the nucleus [25]. Thus, Neat1 can regulate expression of genes transcriptionally by sequestering proteins involved in the regulation of gene promoters, and co-transcriptionally by binding to mRNAs. While some studies have been undertaken to determine the sequence elements in Neat1 responsible for the recruitment of proteins [26], little is known about potential specific elements in paraspeckle mRNA targets except that some mRNA targets contain inverted repeats derived from SINE repeats in their 3’ UTRs [8][4] an element we didn’t find in 3’UTRs of paraspeckle mRNA targets in rat GH4C1 cells. Two main strategies can be used to identify recurring sequence patterns or motifs, namely a global computational motif discovery approach or a gene candidate strategy. The first one adopts either traditional and popular methods such as MEME (Multiple Expression motifs for Motif Elicitation) [27] or more recent one such as Amadeus (A Motif Algorithm for Detecting Enrichment in mUltiple Species) [28] to discover the sequence consensus of RNA Binding Protein’s binding sites or used Z-score statistics to search for the overrepresented nucleotides of a certain size. In the case of our issue, the main practical limitations of these genome-scale detections of known and/or novel motifs lies in the fact that the known or novel motif has to be overrepresented to be identified and the size of the searched sequence has to be defined. However, a nucleotide sequence may be involved in paraspeckle binding without being overrepresented in paraspeckle mRNA targets, like it is the case for IRAlu sequences and the size of the potential sequence cannot be presupposed a priori. For all these reasons, a gene candidate strategy appeared more suitable to reach our goal.

Among the 3928 paraspeckle RNA targets we reported in GH4C1 cells [12], our choice of gene candidate fell on Calr mRNA, which is shown here to be closely associated not only with Neat1 but also with every major core-protein component of paraspeckles. Moreover since a circadian post-transcriptional regulation is shown here to account for its circadian expression pattern as previously shown in the liver [16] and since we previously reported that its circadian nuclear retention is disrupted after Neat1 knock-down [12], it may be assumed that paraspeckle binding is crucial for Calr mRNA expression and circadian pattern [12]. Finally, the 3’UTR of Calr mRNA appears here as efficient as an IRAlu sequence to bind Neat1 and to induce a nuclear retention of a reporter gene, leading to the conclusion that the dissection of 3’UTR Calr mRNA may allow to uncover sequence elements involved in paraspeckle binding.

The most proximal and the most distal regions of the 3’UTR Calr mRNA are further identified as two regions able to associate with Neat1. However in spite of their efficient association with paraspeckles, these two regions are far less efficient than the entire 3’UTR to induce a nuclear retention of the reporter gene. It then seems that each of these regions can bind to paraspeckles with a binding strength that is insufficient to allow for their efficient nuclear retention. In addition to show that Calr mRNA needs to be anchored by at least two sites to be efficiently retained in the nucleus by paraspeckles, more generally, these results suggest that cooperative effects between different parts of 3’UTR may be necessary for an efficient nuclear retention of the mRNA. It is tempting to speculate that these cooperative effects between different parts of the 3’UTR may be sustained by secondary structures carried by mRNA and as previously suggested may be determinant for nuclear retention [29]. In any case, the efficiency of the most proximal and the most distal regions of 3’UTR Calr to associate with paraspeckles prompted us to further analyze their molecular features.

Using RNA-RNA interaction prediction algorithms (IntaRNA, http://rna.informatik.uni-freiburg.de/) [30] we were able to ruled out that the two identified regions of 3’UTR Calr can directly hybridize with Neat1. Indeed, the only part of 3’UTR Calr that offered a significant prediction score for Neat1 interaction is located between the two regions of interest (see Supplemental Fig 3). While devoid of predictable Neat1 direct interaction sequence, the two regions of interest were shown to display after alignment a common sequence of 15 nucleotides with Y in 5^th^ and R in 11^th^ position. These two 15-nucleotide sequences found in 3’UTR of Calr mRNA are shown to participate in paraspeckle binding and Calr mRNA nuclear retention since their mutation reduces both binding and nuclear retention of the reporter gene. Interestingly 30% of paraspeckle mRNA targets arbor a sequence close to this one in their 3’UTR, with an average of 1.6 sequences per 3’UTR. However, the occurrence of this sequence in 3’UTR is not overrepresented in mRNA that are paraspeckle targets as compared to all expressed mRNA in the cells. It then appears that while the 15-nucleotide sequence identified here may allow the 3’UTR of mRNAs to associate with paraspeckles, its occurrence by itself in 3’UTR of mRNAs does not make them paraspeckle targets. As already mentioned, the same conclusion can be drawn for IRAlu sequences. In any case, this 15-nucleotide sequence is shown here to be present twice more frequently than a random sequence of same length in the 3’UTR of all mRNAs. This high prevalence could support an important role of this sequence in RNA biology.

By mass spectrometry analysis, 24 proteins were identified as specifically bound to an oligonucleotide containing the 15-nucleotide sequence we reported here. As expected, a majority of them are RNA binding proteins that form a dense plexus of interactions between each other. Among them, only one, namely HNRNPK, has been previously described as a paraspeckle protein component [25]. Focusing on this protein, it was found that HNRNPK is able to directly bind the oligonucleotide containing the 15-nucleotide sequence identified.

Interestingly, the motif enriched in the HNRNPK binding peaks throughout the transcriptome, as identified by GraphProt [31] has been shown to correspond to a pyrimidine-rich sequence [29], consistent with the known preferences of HNRNPK for C-rich sequences [32]. This appears in accordance with the 15-nucleotide sequence we identified here, in which more than half of the nucleotides are pyrimidines. Moreover, the three KH RNA-binding domains of HNRNPK were previously shown to act cooperatively in binding sequences with triplets of C/T-rich regions [33] fitting the sequence architecture of the entire 3’UTR Calr mRNA. However, while the presence of multiple cytosine patches appears to be necessary for interaction with HNRNPK, it is not always sufficient and does not guarantee an interaction with HNRNPK [34]. This is illustrated here by the inability of the middle region of the 3’UTR of Calr mRNA (3’UTR-C2 fragment) to associate with paraspeckles in spite of numerous cytosine patches (see Supplemental Fig. 3). It has also been suggested that the C/T-rich motif is preferentially bound in a structured context. The interplay of C-patch spacing and secondary structure formation influences HNRNPK RNA recognition since HNRNPK recognition of C-patches depends on positioning within the RNA structure [34]. Interestingly, predictive structural analysis of the 30 bases oligonucleotide we used in the HNRNPK binding experiments, demonstrates the accessibility of C-patches located both inside the 15-nucleotide sequence identified here and in its 5’ adjacent region (Supplemental Fig. 6). It is then tempting to speculate that the binding of HNRNPK involves either the 15-nucleotide sequence or the adjacent 5’ region or both.

A 42 nucleotides fragment that contained three stretches of at least six pyrimidines (C/T) which has been named SIRLOIN (SINE-derived nuclear RNA LOcalizatIoN) has been previously shown to bind HNRNPK and to direct nuclear enrichment of mainly lncRNAs but also some mRNAs [29]. As suggested by these authors and in view of our present results, multiple independent pathways are likely responsible for nuclear enrichment in lncRNAs and mRNAs, recognizing specific RNA sequences and/or other features of the ribonucleoproteins.

The present results allow from the 1894 sequences found in 3’UTR of paraspeckle mRNA targets from a pituitary cell line to design a motif and to propose a novel matrix in 3’UTR of mRNAs that may contribute to their nuclear retention through association with paraspeckles. Furthermore, these results suggest that the recognition of this sequence in a structured RNA context by HNRNPK is important to bridge paraspeckles with their mRNA targets. In this view, HNRNPK appears then not only essential for paraspeckle formation [10] but also for nuclear retention of paraspeckle mRNA target. However the sequence we described here is widely present in 3’UTR of mRNAs whether they are or they aren’t paraspeckle targets and it thus may be assumed that other specific RNA sequences and/or other associated proteins are involved in mRNA paraspeckle nuclear retention. This underlines that description of a simple motif recognition site to predict the complex binding events may be overly simplistic and provide an inaccurate view of binding events that occur through alternate modes of accommodation. Nevertheless, based on these findings, the novel matrix proposed can be presented as a promising candidate for the elusive recruiting elements of mRNAs by paraspeckles and lay the groundwork for future studies to interrogate the role of cooperative other sequences in paraspeckle mRNA nuclear retention.

## SUPPLEMENTARY DATA

Supplemental Table 1: Sequences of qPCR primers and oligonucleotides

Supplemental Table 2: Consensus matrix probabilities built from the 1894 sequences found in 3’UTR of Neat1 mRNA targets

Supplemental Fig. 1: Neat 1 RNA pull-down controls in the different experiments

Supplemental Fig. 2: Dual FISH visualization of Calr mRNA localizing to paraspeckles

Supplemental Fig. 3: Sequence of cloned 3’UTR-Calr with delineation of C1, C2 and C3 fragments

Supplemental Fig. 4: Prevalence of the 15-nucleotide sequence compared to a random sequence Supplemental Fig. 5: Heat map of the 45 quantified proteins in RNA-protein pull-down

Supplemental Fig. 6: Predicted lowest energy structures of the 30 oligonucleotides probe used in RNA-protein pull down as determined by RNAfold web server (http://rna.tbi.univie.ac.at/cgi-bin/RNAWebSuite/RNAfold.cgi)

## DECLARATIONS

### DATA AVAILABILITY

The mass spectrometry proteomics data have been deposited to the ProteomeXchange Consortium via the PRIDE [1] partner repository with the dataset identifier PXD020458.

