## Supplemental Table 1, Fig 1, Fig 2, Fig 3, Fig 4, Fig 5, Fig 6 for "A Sequence determinant in 3’UTR of mRNAs for Nuclear Retention by Paraspeckles"

**Supplemental Table 1: Sequences of qPCR primers and oligonucleotides**

**<u>qPCR Primer list (rat) (5' to 3')</u>**

Neat1-1 and 1-2
AAGGCACGAGTTAGCCGCAAAT
TGTGCACAGTCAGACCTGTCATTC

Neat1-2
GCCTGCTTTCAGCTGTTGGTTT
TCTGGACAGCAACTGAGCAATACG

Egfp
CGGCATCAAGGTGAACTTCAAGATCC
ACTGGGTGCTCAGGTAGTGGTT

Calr pre-mRNA
GTGTCCACCTCTGTTCATCTGGTT
AGAGGTCTAAGCCCAGTACAGCAA

Calr
ACCAGAAGGACATGCATGGAGACT
TTGTTGATCAGCACGTTCTTGCCC

Gapdh
CCCTCAAGATTGTCAGCAATG
GTCCTCAGTGTAGCCCAGGAT

**<u>Calr 3' RACE</u>**

GSP1 : CAGAGAAGCAGATGAAGGACAAGCAG
GSP2 : GCCAAGCCAAGGATGAGCTGT

**Cloning of 3'UTR Calr fragments**

**C1**: GCCAAGCCAAGGATGAGCTGTAG
AATCAGAATCCACCCCAGACCTGAAC

**C2**: TCAGGTCTGGGGTGGATTCTGATTT
CCTAGGGCTTTTCCTCCATACCTGT

**C3**: AGCCCTAGGCTTGAGATTTCATCTGC
ACACTCTCAGTGTGAGCTGTGCTA

**Mutagenesis of C1 and C3**

**$C1_M$ :**

Forward 5'-GAGGCCACACCACCAGGCACGACGCCAGCACTGAGGCCTGAAC-3'

Reverse 5'- GTTCAGGCCTCAGTGCTGGCGTCGTGCCTGGTGGTGTGGCCTC-3'

**$C3_M$ :**

Forward 5'-GCTCTTCCCCTTTCTCCCTAGGCGCGAGGTCAGCGCCATTTGTGGG-3'

Reverse 5'-CCCACAAATGGCGCTGACCTCGCGCCTAGGGAGAAAGGGGAAGAGC-3'

**Neat1 RNA pull-down specific oligonucleotides (3' biotinylated with a triethyleneglycol spacer)**

S oligo 1: CTCCACCATCATCAATCCTCTGGAC

S oligo 2: GCCTTCCCACATTTAAAAACACAAC

Non-specific: ATAATTTCAAACATCAAATGGTATTTTA

**RNA protein pull-down specific oligonucleotides (3' biotinylated with a triethyleneglycol spacer)**

Specific oligonucleotide : U*C*U*C*C*C*UUGCCCCCAGGACUGGGCCAUUUG
*Phosphorothioate bonds

Negative RNA Control [poly(A)$_{25}$ RNA] (PierceTM Magnetic RNA-Protein Pull-Down Kit

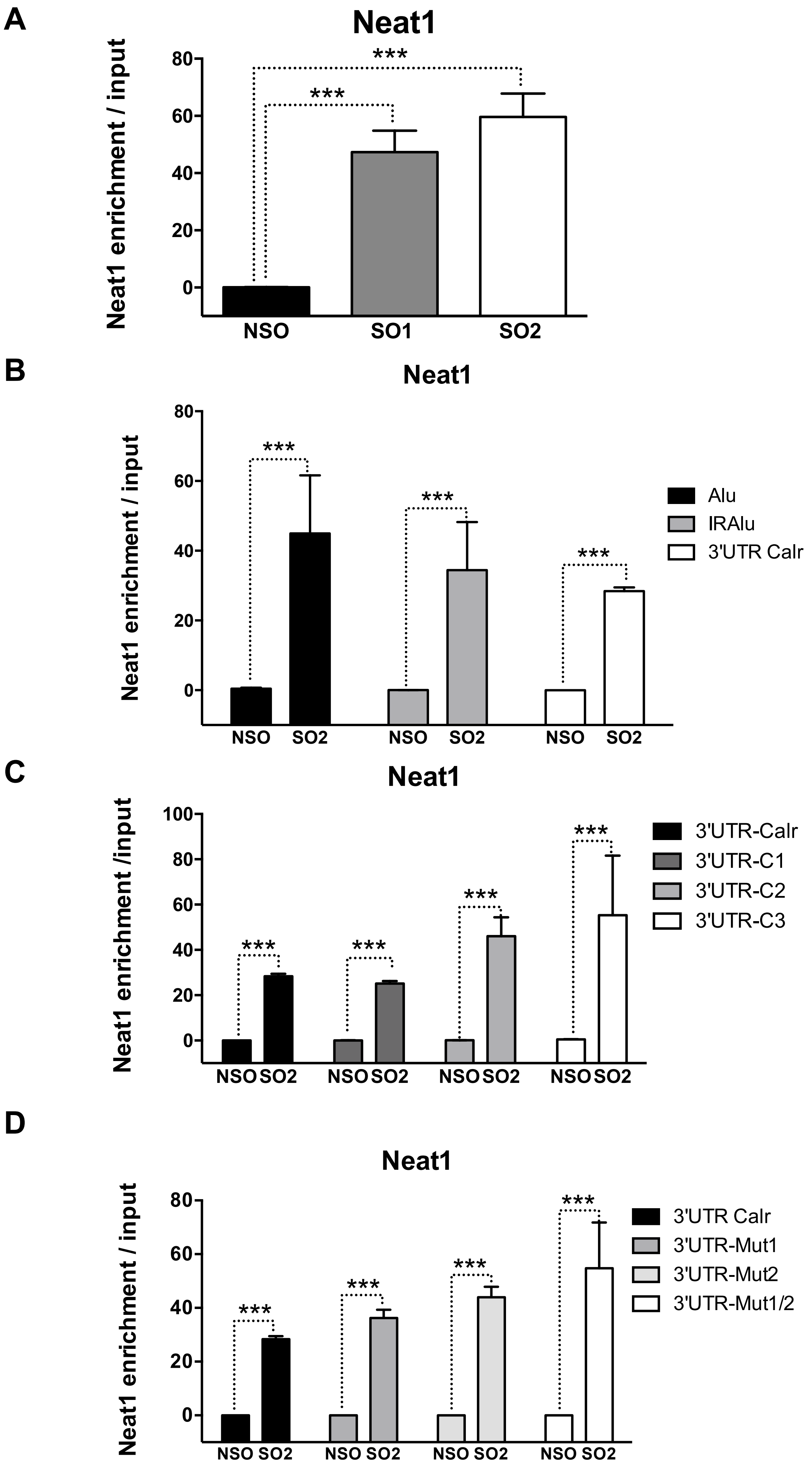
A
Neat1
Neat1 enrichment / input
80
60
40
20
0
***
***
NSO
SO1
SO2
B
Neat1
Neat1 enrichment / input
80
60
40
20
0
***
***
***
Alu
IRAlu
3'UTR Calr
NSO
SO2
NSO
SO2
NSO
SO2
C
Neat1
Neat1 enrichment /input
100
80
60
40
20
0
***
***
***
***
3'UTR-Calr
3'UTR-C1
3'UTR-C2
3'UTR-C3
NSO SO2
NSO SO2
NSO SO2
NSO SO2
D
Neat1
Neat1 enrichment / input
80
60
40
20
0
***
***
***
***
3'UTR Calr
3'UTR-Mut1
3'UTR-Mut2
3'UTR-Mut1/2
NSO SO2
NSO SO2
NSO SO2
NSO SO2

**Supplemental Fig. 1: Neat 1 RNA pull-down controls in the different experiments**

**A.** Enrichment of Neat1 RNA relative to input after RNA pull-down in native GH4C1 cells. Enrichment obtained after using the two anti-sens specific (Specific Oligonucleotide 1: SO1 and Specific Oligonucleotide 2 : SO2) or a non-specific (NSO) oligonucleotide probes. ***$p<0.001$vs non-specific oligonucleotide probe. **B.** Enrichment of Neat1 RNA relative to input after RNA pull-down in Alu-, IRAlu- and 3'UTR-Calr-containing Egfp mRNA cell lines. Enrichment obtained after using the anti-sens specific (Specific Oligonucleotide 2 : SO2) or a non-specific (NSO) oligonucleotide probes. ***$p<0.001$vs non-specific oligonucleotide probe. **C.** Enrichment of Neat1 RNA relative to input after RNA pull-down in 3'UTR-Calr-, 3'UTR-C1-, 3'UTR-C2- and 3'UTR-C3-containing Egfp mRNA cell lines. Enrichment obtained after using the anti-sens specific (Specific Oligonucleotide 2 : SO2) or a non-specific (NSO) oligonucleotide probes. ***$p<0.001$vs non-specific oligonucleotide probe. **D.** Enrichment of Neat1 RNA relative to input after RNA pull-down in 3'UTR-Calr-, 3'UTR-Mut1-, 3'UTR-Mut2- and 3'UTR-Mut1/2-containing Egfp mRNA cell lines. Enrichment obtained after using the anti-sens specific (Specific Oligonucleotide 2 : SO2) or a non-specific (NSO) oligonucleotide probes. ***$p<0.001$vs non-specific oligonucleotide probe.

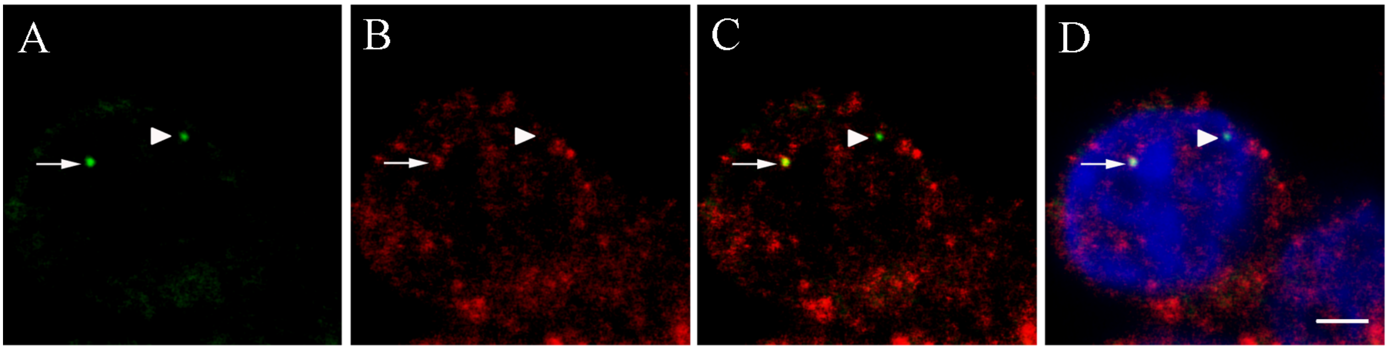

Neat1 Calr mRNA merged merged+ Dapi

**Supplemental Fig. 2: Dual FISH visualization of Calr mRNA localizing to paraspeckles**
Dual Calr and Neat1 RNA-FISH to pituitary GH4C1 cells show **A.** the nuclear distribution of Neat1 RNA in a few distinct foci (arrow and arrow head). **B.** the cytoplasmic and diffuse nuclear localization of Calr mRNA and its distribution in some nuclear distinct foci. **C.** foci in which Calr and Neat1 RNA overlap (arrow) indicating the paraspeckle localization of Calr mRNA and foci without overlap (arrow head). **D.** Nuclear staining by Hoechst is added to C. Scale bars equal 5 μm.

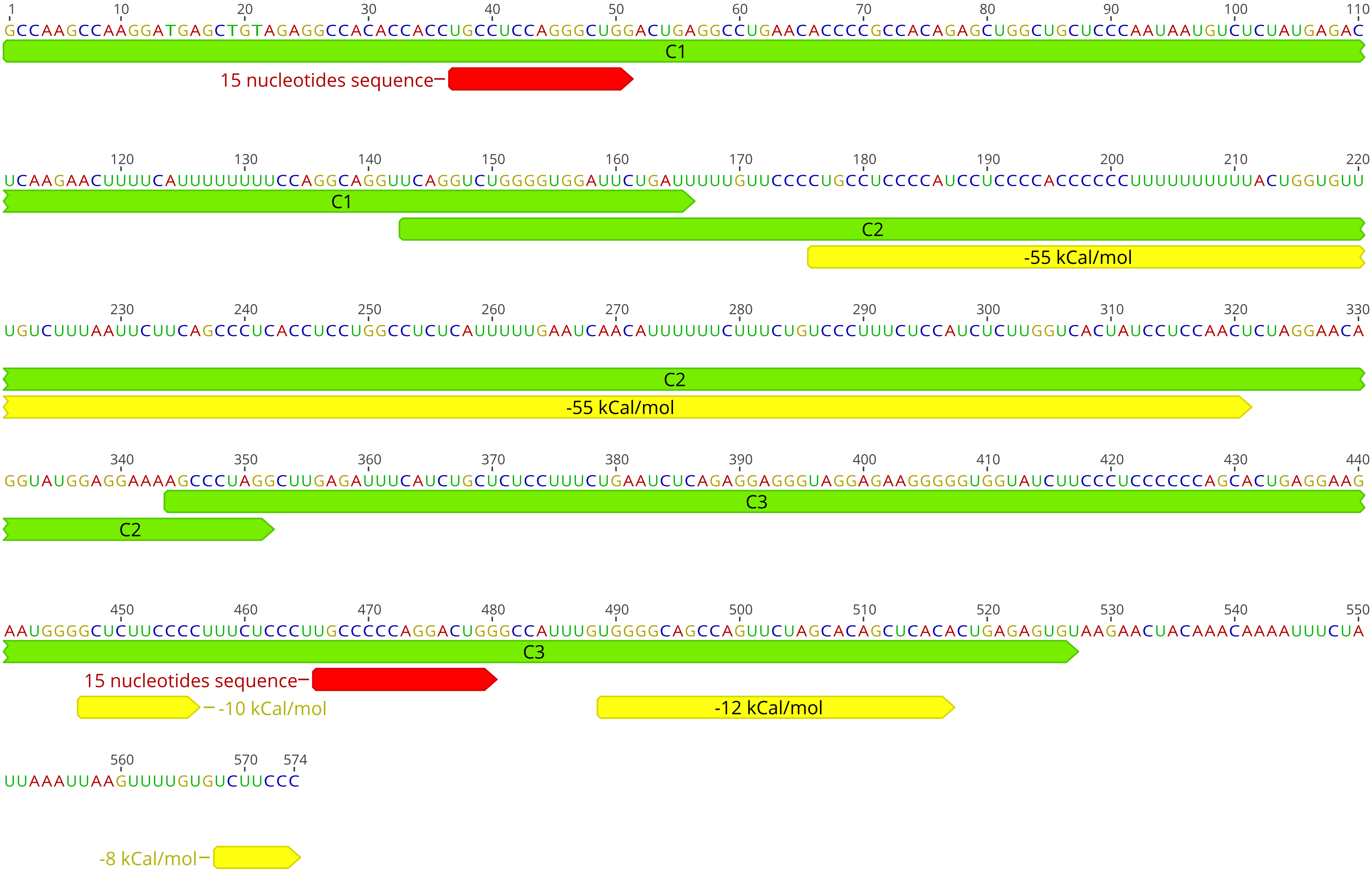

1 10 20 30 40 50 60 70 80 90 100 110
GCCAAGCCAAGGATGAGCTGTAGAGGCCACACCACCUGCCUCCAGGGCUGGACUGAGGCCUGAACACCCCGCCACAGAGCUGGCUGCUCCCAAUAAUGUCUCUAUGAGAC
C1
15 nucleotides sequence
120 130 140 150 160 170 180 190 200 210 220
UCAAGAACUUUUCAUUUUUUUUCCAGGCAGGUUCAGGUCUGGGGUGGAUUCUGAUUUUUGUUCCCCUGCCUCCCCAUCCUCCCCACCCCCCUUUUUUUUUACUGGUGUU
C1
C2
-55 kCal/mol
230 240 250 260 270 280 290 300 310 320 330
UGUCUUUAAUUCUUCAGCCCUCACCUCCUGGCCUCUCAUUUUUGAAUCAACAUUUUUUCUUUCUGUCCCUUUCUCCAUCUCUUGGUCACUAUCCUCCAACUCUAGGAACA
C2
-55 kCal/mol
340 350 360 370 380 390 400 410 420 430 440
GGUAUGGAGGAAAAGCCCUAGGCUUGAGAUUUCAUCUGCUCUCCUUUCUGAAUCUCAGAGGAGGGUAGGAGAAGGGGGUGGUAUCUUCCCUCCCCCCAGCACUGAGGAAG
C3
C2
450 460 470 480 490 500 510 520 530 540 550
AAUGGGGCUCUUCCCCUUUCUCCCUUGCCCCCAGGACUGGGCCAUUUGUGGGGCAGCCAGUUCUAGCACAGCUCACACUGAGAGUGUAAGAACUACAAACAAAAUUUCUA
C3
15 nucleotides sequence
-10 kCal/mol
-12 kCal/mol
560 570 574
UUAAAUUAAGUUUUGUGUCUUCCC
-8 kCal/mol

**Supplemental Fig. 3: Sequence of cloned 3'UTR-Calr with delineation of C1, C2 and C3 fragments (in green).** The 15-nucleotide sequences (in red) found in C1 and C3 are positioned. Regions in 3'UTR exhibiting a significant prediction score for Neat1 interaction are indicated in yellow. The most significant prediction score is found in C2 (-55kCal/mol).

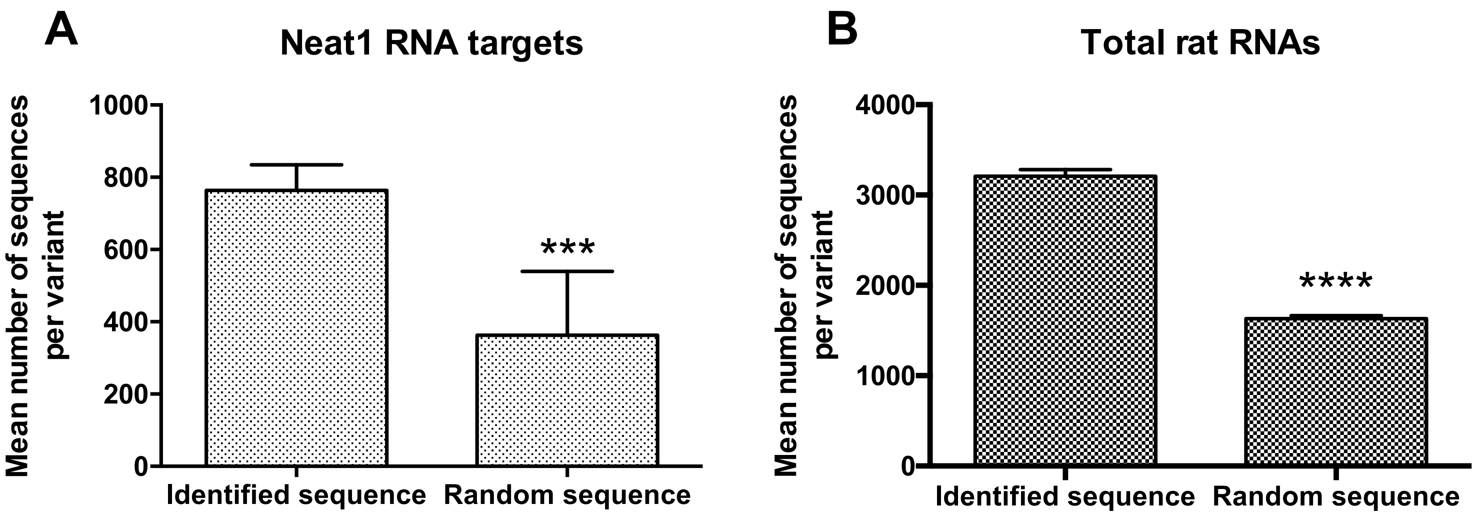

A
Neat1 RNA targets
Mean number of sequences per variant
1000
800
600
400
200
0
***
Identified sequence
Random sequence
B
Total rat RNAs
Mean number of sequences per variant
4000
3000
2000
1000
0
****
Identified sequence
Random sequence

**Supplemental Fig.4: Comparison of the number of occurrences of the 15-nucleotide sequence identified here with that of a 15-nucleotide sequence randomly generated, both with Y or R in 5$^{th}$ and 11$^{th}$ position. A.** Mean values for the 4 variants of the 15-nucleotide sequence and for the 4 variants of 30 randomly generated 15-nucleotide sequences found in the 3'UTR of Neat1 RNA targets**. B.** Mean values for the 4 variants of the 15-nucleotide sequence and for the 4 variants of 30 randomly generated 15-nucleotide sequences found in the 3'UTR of all rat RNAs**.** ***p<0.01 ****p<0.001

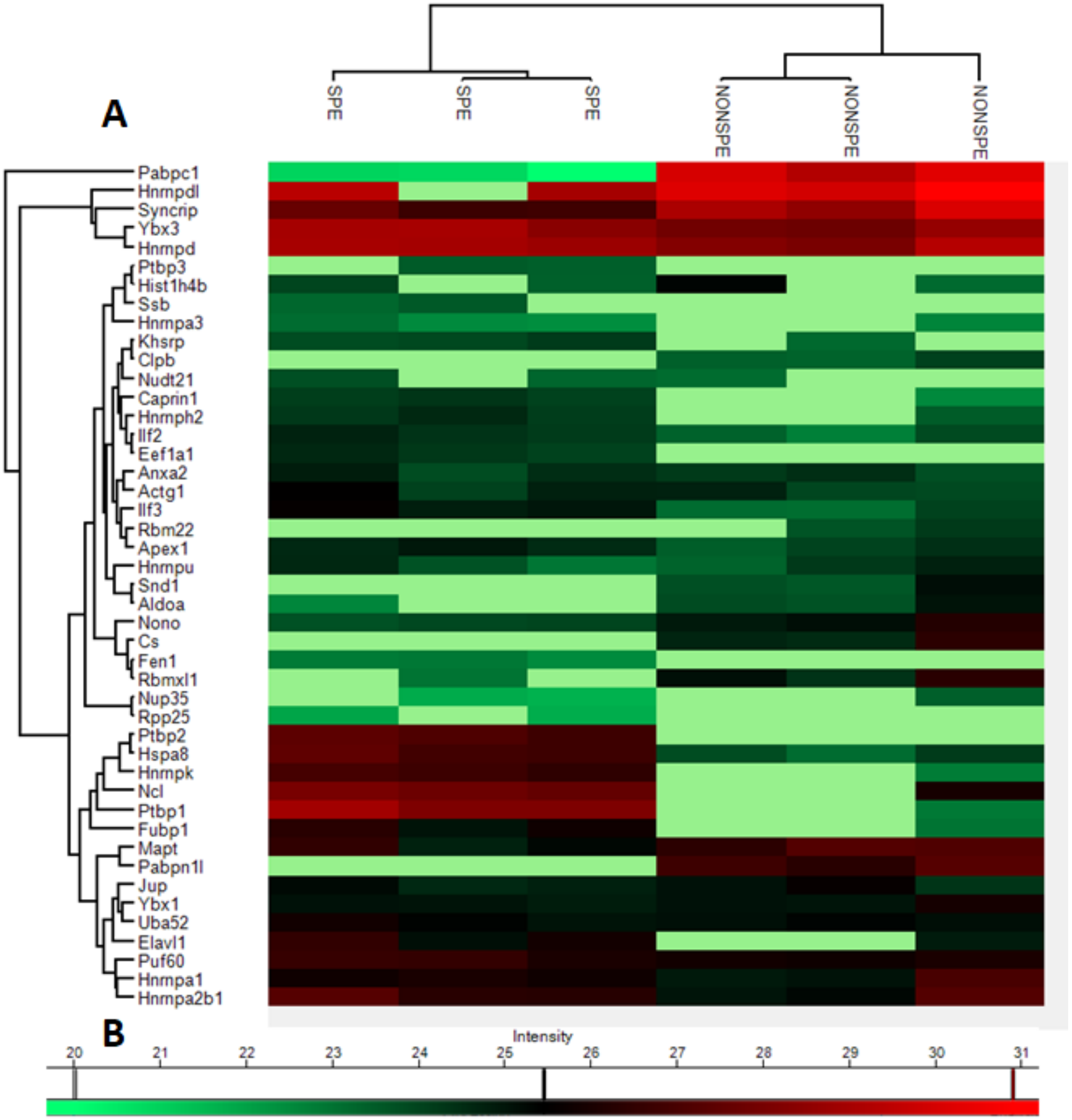
A
SPE
SPE
SPE
NONSPE
NONSPE
NONSPE
Pabpc1
Hnrnpdl
Syncrip
Ybx3
Hnrnpd
Ptbp3
Hist1h4b
Ssb
Hnrnpa3
Khsrp
Clpb
Nudt21
Caprin1
Hnrnph2
Ilf2
Eef1a1
Anxa2
Actg1
Ilf3
Rbm22
Apex1
Hnrnpu
Snd1
Aldoa
Nono
Cs
Fen1
Rbmxl1
Nup35
Rpp25
Ptbp2
Hspa8
Hnrnpk
Ncl
Ptbp1
Fubp1
Mapt
Pabpn1l
Jup
Ybx1
Uba52
Elavl1
Puf60
Hnrnpa1
Hnrnpa2b1
B
Intensity

**Supplemental Fig. 5: (A)** Heat map of the 45 quantified proteins in RNA-protein pull-down The proteins are designated by their gene symbol, SPE means binding to biotinylated RNA containing the 15-nucleotide sequence and NONSPE is for non-specific probe (Negative RNA Control [poly(A)$_{25}$ RNA] from the PierceTM Magnetic RNA-Protein Pull-Down Kit). Three replicates using the specific probe and three replicates using a non-specific probe are shown. The light green blocks correspond to the missing values without imputation. **(B)** Log intensity scale of protein intensities.

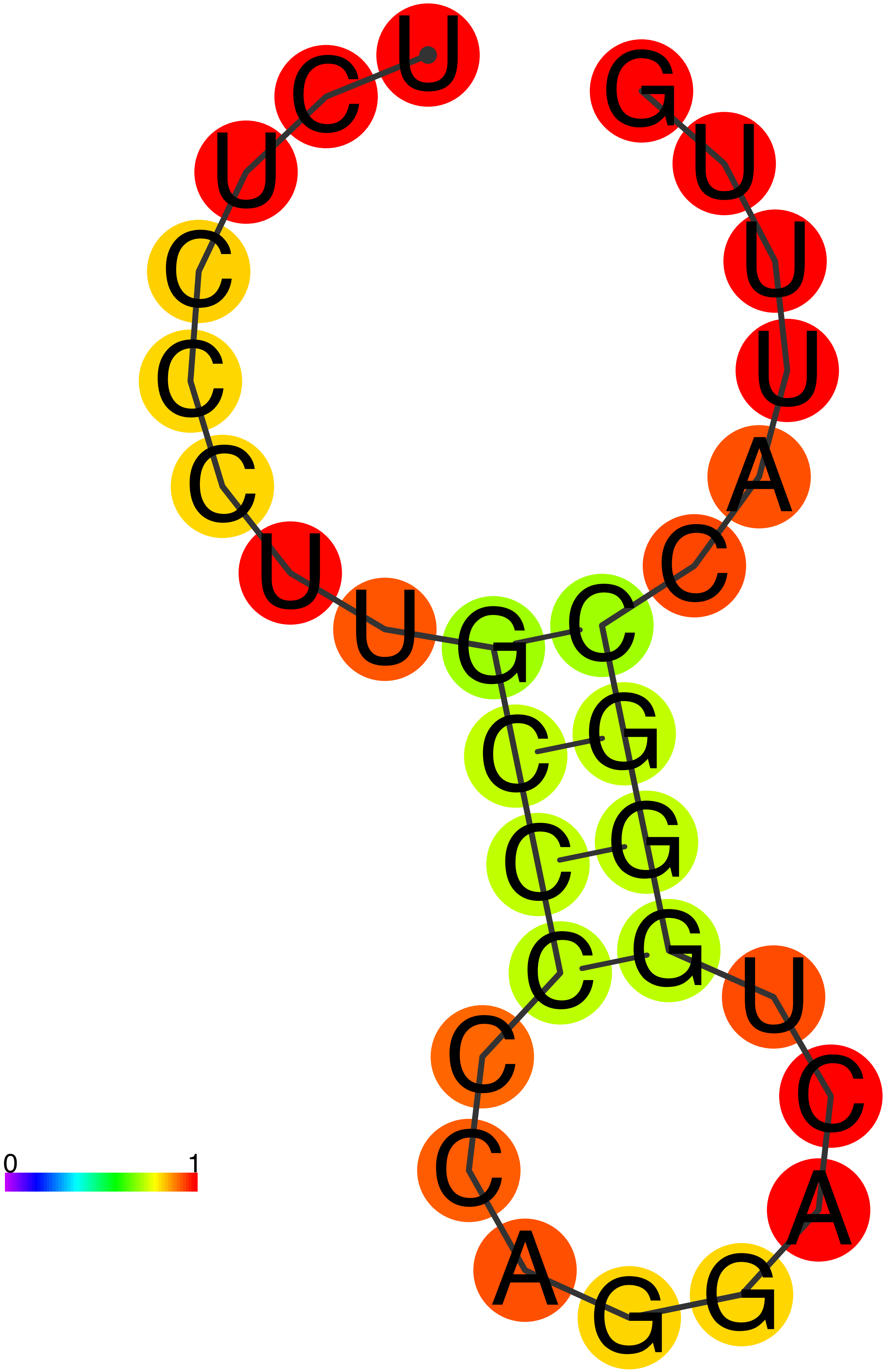
0
1

**Supplemental Fig. 6:** Predicted structures of the single stranded 30-nucleotide RNA sequence by RNAfold web server (http://rna.tbi.univie.ac.at/cgi-bin/RNAWebSuite/RNAfold.cgi). Structure is based on the minimal free energy (MFE) method (an established method to predict RNA structure). Complimentary regions are evaluated to predict the most energetically stable molecule.
